# Diatom biodiversity response to shrinking glaciers in the Peruvian Andes

**DOI:** 10.1101/2025.02.17.635084

**Authors:** Nicky R. Kerr, Duncan Quincey, Sarah C. Fell, Martyn Kelly, Katy Medina, Edwin Loarte, Lee E. Brown

**Author notes:** Corresponding author: Nicky R. Kerr.

## Abstract

Alpine river biodiversity is under threat as climate change is driving extensive glacier retreat worldwide, altering aquatic habitats and restructuring biotic communities. However, with the exception of macroinvertebrates, the responses of aquatic groups to deglaciation have received little attention, particularly in tropical ecosystems where species may respond differently owing to additional pressures such as distinct wet and dry seasons, limited snowmelt and extreme high altitude and thus lower oxygen availability. Utilising a continuum of glacier cover across 23 rivers in the Peruvian Andes (Cordilleras Blanca and Vilcanota), we demonstrate that diatoms, abundant primary producers in glacier-fed rivers, show regionally consistent increased α-diversity and density responses with deglaciation. However, β-diversity was found to decline, potentially reflecting the reduction of habitat heterogeneity and niche space both within and between alpine rivers as glaciers are lost. High regional biodiversity, with just 23% of taxa found in both regions, was closely linked to the distinct underlying geologies of the Andes which influence water chemistry. Thirty-eight taxa were found exclusively at sites with glacier cover ≥ 25%, which is considerably higher than numbers recorded in European alpine rivers. However, 31 taxa were identified only at non-glacial sites suggesting deglaciation could open up more habitat for these species. Peruvian rivers overall showed similar responses to glacier retreat as temperate alpine systems, but the large number of taxa found only at mid-high glacial influence sites suggests tropical diatom assemblages may be especially vulnerable to climate change.

## Introduction

Glaciers currently cover approximately 10% of Earth’s land surface and store about 50% of the world’s freshwater supply but they are shrinking rapidly across the world (IPCC, 2019; Rounce et al., 2023). In tropical regions like the Peruvian Andes, this poses critical risks to freshwater ecosystems that sustain biodiversity, nutrient cycling and essential water supplies for local communities (Encalada et al., 2019). Over 70% of the world’s tropical glaciers are found in Peru and the Andes are characterised by extreme seasonality and high altitude stressors (Seehaus et al., 2019). However, warming and more varied precipitation patterns have forced rapid glacier retreat for several decades, and approximately 69 (± 25) % of remaining glacier ice mass in low latitude regions, such as the Peruvian Andes, is projected to be lost by 2100 (Rounce et al., 2023). Glacier retreat alters the timing and magnitude of meltwater generation and changes the proportional input of other water sources (e.g. groundwater) (Brown et al., 2009). This subsequently alters river habitat conditions such as water temperature, nutrient content and suspended sediment which, in turn, drives changes to biological assemblages (Füreder et al., 2001; Milner et al., 2017). Research seeking to understand how river ecosystems respond to glacier retreat has focused predominantly on macroinvertebrates and fish, with important primary producer groups such as diatoms remaining understudied (Fell et al., 2017).

Diatoms are hard-bodied algae common in all river ecosystems which form the base of aquatic food webs and play major roles in nutrient cycling (Guo et al., 2016). Diatoms are highly sensitive to environmental change, with communities restructuring rapidly as conditions alter (Kelly et al., 2008). Outside of Europe and North America, research on the impacts of glacier retreat on diatoms has been sparse, despite the potential loss of diatom biodiversity shown in temperate regions as glaciers recede (Hieber et al., 2001; Hansen et al., 2006; Fell et al., 2018; Brahney et al., 2021). Riparian-based organic matter input to alpine rivers is minimal (Clitherow et al., 2013; Fellman et al., 2015), and in combination with the Andes’ extreme seasonality and glacier dynamics, understanding diatom assemblage change is critical to assess climate change impacts on river ecosystem functioning (Kohler et al., 2024).

Glacier-fed rivers are known to host different diatom assemblages to non-glacierised sites (Hieber et al., 2001; Bona et al., 2012; Brahney et al., 2021). Across several regions, taxa such as *Hannaea arcus*, a cold water specialist, and *Achnanthidium minutissimum*, known for withstanding harsh environmental conditions, are characteristic of glacier fed rivers (Cantonati, 2001; Hansen et al., 2006; Fell et al., 2021; Brahney et al., 2021; Kahlert, Rühland, et al., 2022; Bert et al., 2024). A study in the Austrian Alps showed that several taxa were recorded only in rivers with a high percentage of catchment glacier cover, including *Fragilariforma constricta* which is listed as ‘threatened’ on the Red List of Algae for Germany (Fell et al., 2018). Whilst none of the species found exclusively in highly glacierised sites are considered glacial-river specialist taxa (Lange-Bertalot et al., 2017) the identification of threatened taxa, coupled with the geographic and genetic isolation of alpine freshwater environments (Dong et al., 2016), highlights the need to understand how rare or vulnerable taxa within poorly studied alpine regions such as the Peruvian Andes will be impacted as glaciers retreat.

Research from Ecuador showed that algal biomass increased significantly when glacier catchment cover declined to less than 11% (Cauvy-Fraunié et al., 2016); however, diatom biodiversity response was not quantified and, more broadly, knowledge of South American river diatoms is restricted to non-glacial rivers (Morales and Vis, 2007; Benito et al., 2018; Benito and Fritz, 2020). Recent research on macroinvertebrates in Peru found α-diversity to increase and β-diversity to decrease in response to glacier shrinkage (Palacios-Robles et al., 2024), similar to diatom responses found in Austrian rivers (Fell et al., 2018). This suggests abiotic (e.g. changes to physical river habitat) and biotic (e.g. dispersal limitation due to traits) mechanisms may structure glacial river communities similarly across multiple trophic levels, although broad scale spatial co-variables such as geology may also influence diatom assemblages (Benito et al., 2018). Considering the rapid glacier retreat occurring in Peru and additional pressures found in the Andes such as long dry seasons and a lack of snow cover (Vuille et al., 2018), analysing the response of riverine diatoms is needed to fully understand the impacts of climate change on tropical alpine biodiversity. This is particularly important given the role of aquatic producers supporting wider food webs, which contribute to the provision of food and clean water to many remote communities within the Andes (Madrigal-Martínez et al., 2022).

This research aims to advance our understanding of community composition and biodiversity of epilithic river diatoms to changes in glacier cover in two alpine regions of the Peruvian Andes: Cordillera Blanca and Cordillera Vilcanota. By studying two discrete mountain areas, diatom responses to a common driver (glacier shrinkage) can be understood in the context of regional differences such as geology, altitude and hydroclimate. The study aims to test the following hypotheses (1) declining glacier cover, associated with the amelioration of harsh habitat conditions, will result in increased α-diversity and abundance of epilithic diatom communities, (2) diatom β-diversity both within and between sites will decrease with glacier shrinkage as river habitats become more similar, and (3) taxon-level responses to declining glacier cover will vary both within and between regions, with some taxa increasing in abundance, whilst others disappear, as glacial meltwater input decreases.

## Methods

### Study areas

Glaciers within the Cordillera Blanca and Cordillera Vilcanota are in long-term retreat, with nearly 25% of the glacierised area in Cordillera Blanca lost since the end of the 20^th^ century (Motschmann et al., 2020) and over 50% of ice area lost since the 1970s in Cordillera Vilcanota (Taylor et al., 2022). The geology of Cordillera Blanca is composed predominantly of granodiorite, an intrusive igneous rock that is part of the Cordillera Blanca batholith (Petford and Atherton, 1992). Cordillera Blanca granodiorite is characterised by quartz, plagioclase feldspar and muscovite mica (Atherton and Sanderson, 1987). Approximately 1800 km south, the geology of Cordillera Vilcanota is more variable with the Chumpe area comprising granitic and sedimentary siliciclastic rock, and the Quelccaya area is volcanic sedimentary (Gomez et al., 2019).

Samples were collected from river sites during September 2019 (Vilcanota) and March 2020 (Blanca) across a gradient of glacier cover in the catchment (GCC), acting as a proxy for different levels of glacier retreat (Figure 1). In Cordillera Blanca, eleven sites were sampled spanning 0-66% GCC within the Parón, Huaytapallana and Llanganuco valleys which drain numerous mountain glaciers. In Cordillera Vilcanota, twelve sites were sampled (0-96% GCC) – eight to the north of Laguna Sibinacocha (Chumpe glacier area), draining from several mountain glaciers and the surrounding catchment and four to the south, draining from the Quelccaya Ice Cap and nearby groundwater seeps (Table 1). River sites were fed by a range of sources, from entirely groundwater (0% GCC; 4x sites) or predominantly groundwater (1-25% GCC; 9x sites), to a combination of groundwater and glacier melt (26-50% GCC; 4x sites) and predominantly glacier melt (> 50% GCC; 6x sites) which provided a range of river characteristics across much of the glacier cover gradient. All sites were situated above the treeline where shading effects were minimal.

**Figure 1.**
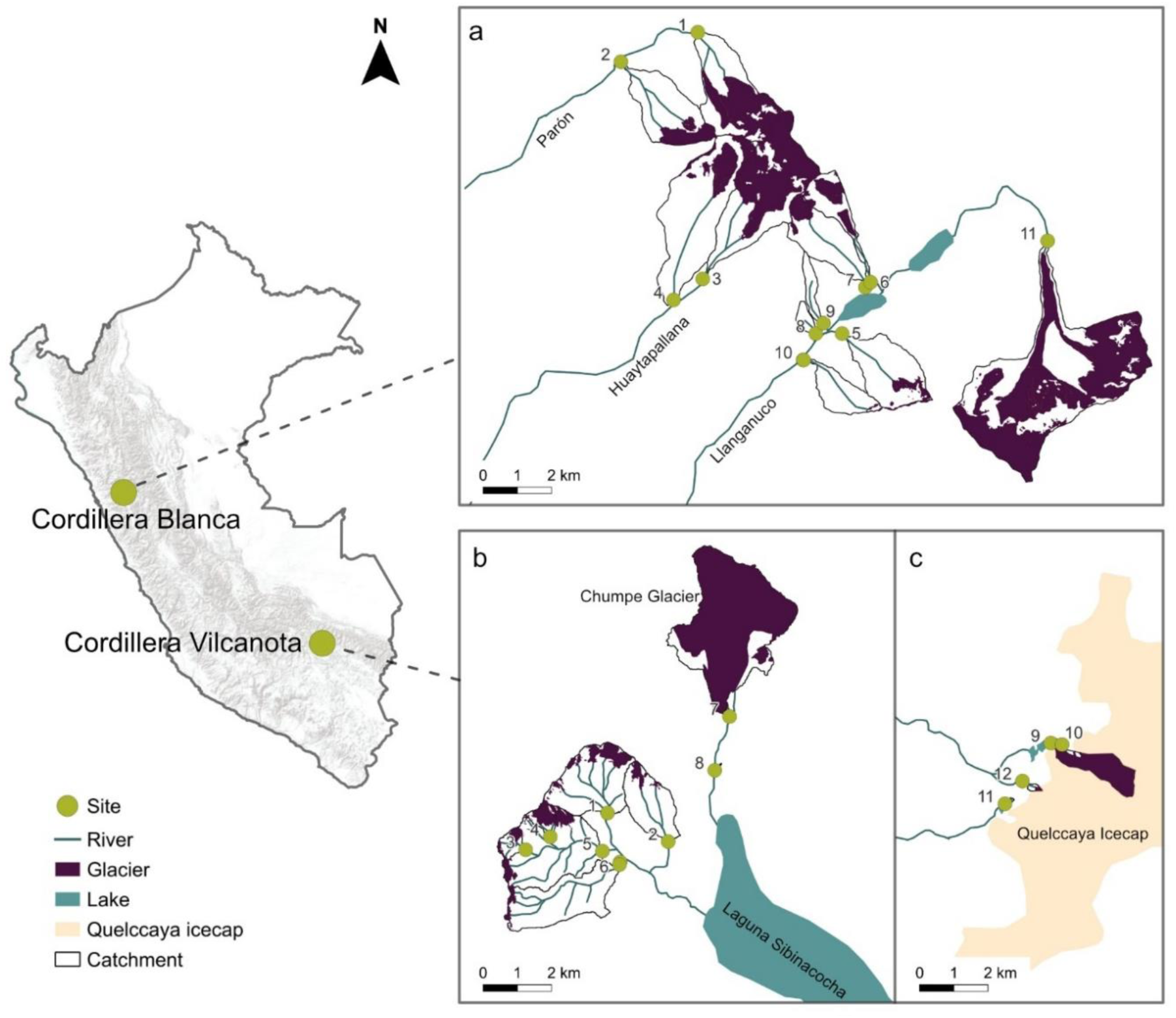
Maps of Peruvian river sites, glaciers and catchments in (a) Cordillera Blanca, (b) Chumpe area of Cordillera Vilcanota and (c) Quelccaya area of Cordillera Vilcanota. The map of Peru (left) is an ESRI terrain map (grey = higher elevation) and shows the location of each Cordillera.

**Table 1.** Sample site locations and information. Glacier cover is the percent of the catchment covered in permanent ice.

| Region | Geology | Catchment | Site number | Altitude (m) | Glacier cover (%) |
| --- | --- | --- | --- | --- | --- |
| **Vilcanota** | Siliciclastic sedimentary | Chumpe | 1 | 5055 | 18 |
|  |  |  | 2 | 5031 | 9 |
|  |  |  | 3 | 5134 | 34 |
|  |  |  | 4 | 5117 | 64 |
|  |  |  | 5 | 5028 | 16 |
|  |  |  | 6 | 5019 | 3 |
|  |  |  | 7 | 4987 | 88 |
|  |  |  | 8 | 4911 | 0 |
|  | Volcanic sedimentary | Quelccaya | 9 | 5191 | 96 |
|  |  |  | 10 | 5204 | 29 |
|  |  |  | 11 | 5179 | 0 |
|  |  |  | 12 | 5211 | 30 |
| **Blanca** | Granodiorite | Parón | 1 | 4442 | 49 |
|  |  |  | 2 | 4218 | 23 |
|  | Granodiorite | Huaytapallana | 3 | 3897 | 62 |
|  |  |  | 4 | 3701 | 19 |
|  | Granodiorite | Llanganuco | 5 | 3888 | 11 |
|  |  |  | 6 | 3908 | 43 |
|  |  |  | 7 | 3822 | 15 |
|  |  |  | 8 | 3805 | 0 |
|  |  |  | 9 | 3793 | 0 |
|  |  |  | 10 | 3666 | 2 |
|  |  |  | 11 | 4198 | 66 |

### Glacier cover

Catchment area upstream of each river site was calculated in QGIS 3.16 using the ‘Channel network and drainage basins’ and ‘Upslope’ functions. To determine water flow direction based on topography, an ALOS PALSAR digital elevation model (DEM; 12.5 m resolution) was used in Cordillera Blanca and a TanDEM-X (10 m resolution) was used in the Cordillera Vilcanota, which were the highest resolution DEMs available. Catchment boundaries were calculated automatically from this. Glacier area in each catchment was determined using a combination of the Normalised Difference Snow Index (NDSI) using Sentinel-2 images, and manual delineation with Sentinel-2, Google Earth imagery and Glacier Land Ice Measurements from Space (GLIMS) polygons (GLIMS Consortium, 2005). Glacier cover in the catchment was then calculated as a percentage of the catchment upstream of the river site that was permanently covered in ice.

### River physical and chemical properties

At each site, measurements of water temperature, electrical conductivity (EC) and pH were taken in Vilcanota using a Hach (Loveland, CO, USA) HQ40D probe, and in Blanca a Xylem (Washington D.C., USA) Multi 3630 multiparameter probe was used. Turbidity was measured with a Cole-Parmer (Vernon Hills, IL, USA) Oakton T-100 turbidity probe. Channel stability was determined using the “channel bottom” component of the Pfankuch Index (Pfankuch, 1975), and then calculating the inverse of this score so that higher scores represented more stable river beds. Water samples were collected by filtering 100 mL of river water through a 0.45 micron filter to determine total nitrogen (TN) and total phosphorus (TP) (Skalar (Breda, Netherlands) San++ automatic wet chemistry analyser) and dissolved inorganic carbon (DIC) and dissolved organic carbon (DOC) (Analytik Jena (Jena, Germany) Multi NC2100 micro elemental analyser) (Table S1).

Benthic diatoms were collected following the CEN water quality and diatom preparation sampling protocol (Comité Européen de Normalisation (CEN), 2014) as applied by Fell et al. (2018) in glacier-fed rivers of Austria. At each site, five submerged cobbles were selected randomly from riffles to represent a diversity of flow and sediment microhabitats (total n = 105 replicates). Using a sterile toothbrush, the benthic biofilm on the upper surface of each cobble was scrubbed from a 9 cm^2^ area using a miniature quadrat. Samples were preserved with Lugol’s iodine prior to identification.

### Diatom identification

To enable the unobstructed observation of diatom valves during microscopic identification, organic material was removed using the hot hydrogen peroxide (H_2_O_2_) method (CEN, 2014) and replicate samples mounted on individual slides. Diatom valves were identified and counted using light microscopy at x 1000 magnification using phase-contrast illumination (Olympus CX43 miscroscope). A minimum of 500 diatom valves were identified from each replicate following multiple taxonomic keys and papers (Kelly, 2000; Krammer and Lange-Bertalot, 2004; Kelly et al., 2005; Krammer and Lange-Bertalot, 2007b; Krammer and Lange-Bertalot, 2007a; Lange-Bertalot et al., 2017; Spaulding et al., 2021) with taxonomic nomenclature following Lange-Bertalot et al. 2017. Where less than 500 valves were present, all valves were counted.

To determine estimates of valve density, the number of valves identified within coverslip transects was used to estimate the total number present on the whole coverslip (0.5 mL) and then multiplied to m^-2^ based on sample volume and rock area (9 cm^2^) sampled (Scott et al., 2014; Fell et al., 2018).

### Data analyses

All data analyses were conducted in R version 4.4.0. A Principal Components Analysis (PCA; *stats* package) was performed on the physical and chemical variables obtained from all sites, to identify underlying patterns within the environmental data and highlight dominant characteristics influencing river sites. Data were first normalised to ensure variables that have different units and measurement scales were weighted equally and the PCA loadings and scores were extracted. Correlations between PC scores with percent glacier cover in the catchment were determined using Pearson’s coefficient.

Biodiversity summary metrics including taxonomic richness, Shannon diversity, Pielou’s evenness and diatom valve density were computed using the *vegan* package (Oksanen et al., 2024). The Shannon diversity index was utilised as many diatom taxa were only identified at single sites in low abundance and it does not weight common taxa higher than rare ones (Morris et al., 2014). The average of each biodiversity metric was then calculated from the five replicates per site. Relationships with GCC were analysed both across and within regions using generalised additive mixed models (GAMM; *mgcv* package (Wood, 2023)), incorporating geology and catchment as random variables.

Within-site β-diversity components total dissimilarity (Sørensen), turnover (Simpson) and nestedness were calculated with the *betapart* package (Baselga et al., 2023) using the valve abundance of taxa identified in individual replicates from each site. Between-site β-diversity was determined first by calculating average taxon abundances across replicates, to result in a single abundance value per taxon per site. Total dissimilarity, turnover and nestedness were then calculated from this taxon-abundance matrix. GAMMs were used to analyse the relationship between GCC and each within-site β-diversity component. Differences in catchment glacier cover between sites within each region were calculated and the relationship between these dissimilarities and each of the between-site β-diversity components was analysed using Mantel tests.

At both regions collectively, diatom community composition and taxon-level responses to decreasing glacier cover were analysed using non-metric multidimensional scaling (NMDS). The average valve abundance of individual taxa at each site was log_10_ (x + 1) transformed to limit the effect of the large maximum abundance data on zero values (Khamis et al., 2014; Fell et al., 2018). The NMDS was applied to a Bray-Curtis dissimilarity matrix of the taxon-abundance data using the *vegan* package. The *envfit* function was used to analyse how the PC1 and PC2 scores (representing river physical and chemical data) and GCC were correlated to the NMDS. To analyse the association between individual species presence and glacier cover levels (None = 0%, Low = 1-25%, Moderate = 26-50%, High = > 50%) an indicator species analysis was performed using the *multipatt* function in the *indicspecies* package (Cáceres et al., 2024).

## Results

### River physical and chemical properties

The PC1 axis accounted for 36% of the variation in the river physical and chemical data and was most closely associated with water chemistry variables including dissolved inorganic carbon (DIC) and electrical conductivity (Figure 2; Table 2). These variables were both higher at sites in the Chumpe area (Cordillera Vilcanota), which was underlain by siliclastic sedimentary rock, leading to a clear separation of catchments along axis 1. The PC2 axis (19% of variation) was most closely associated with physical habitat characteristics such as turbidity and water temperature, along with pH. PC2 scores increased significantly with increased glacier cover in the catchment (R^2^ = 0.29, p = 0.008). The PC1 axis had no significant correlation with glacier cover (Figure S1; Table 2).

**Figure 2.**
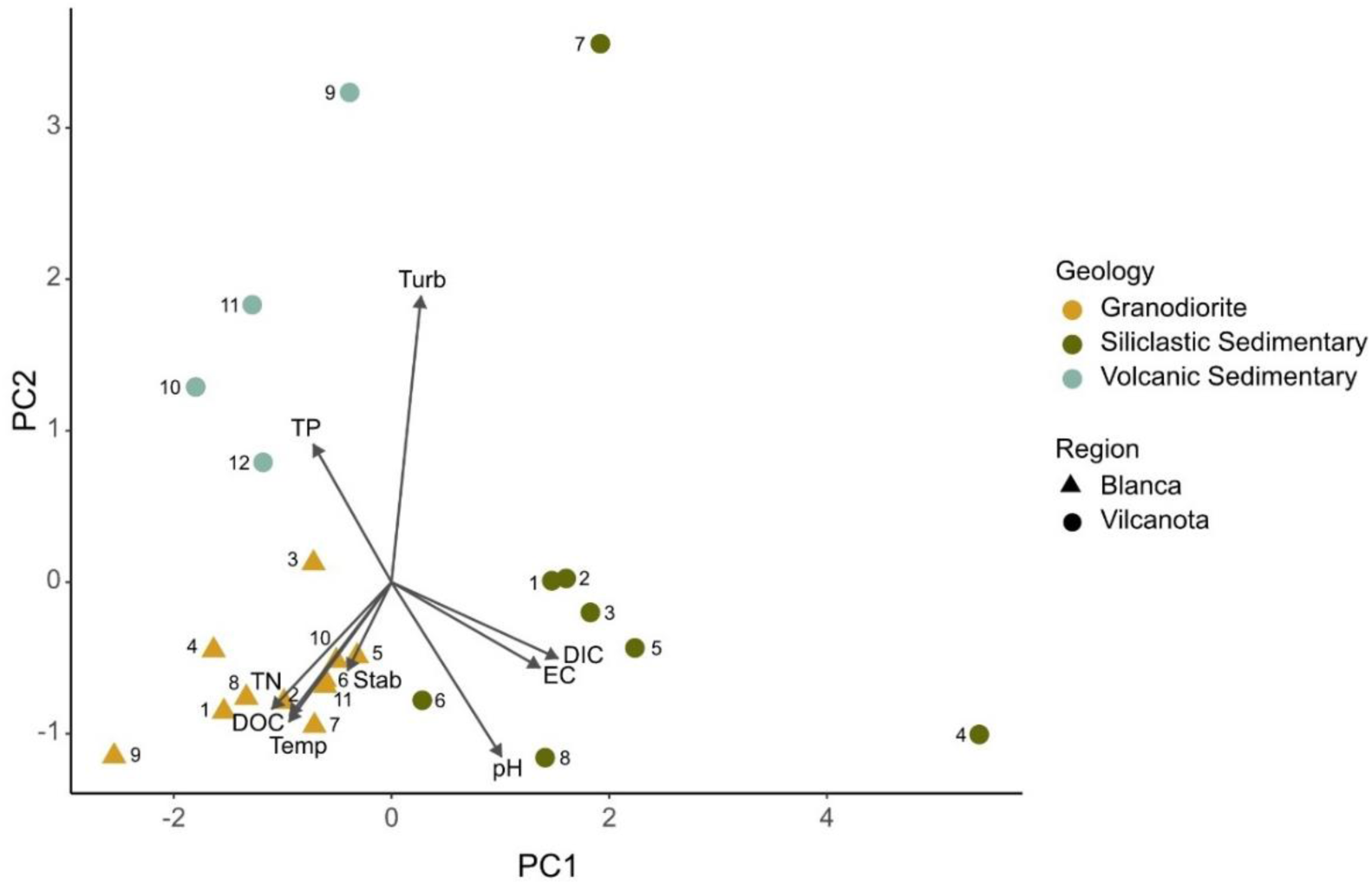
PCA plot of environmental variables from sites in the Cordillera Blanca (triangles) and Cordillera Vilcanota (circles). Numbers refer to the site number for that region. Abbreviations: DIC = dissolved inorganic carbon (mg L^-1^), DOC = dissolved organic carbon (mg L^-1^), EC = electrical conductivity (μS cm^-1^), TP = total phosphorous (mg L^-1^), TN = total nitrogen (mg L^-1^), Stab = channel stability (1/Pfankuch), Temp = water temperature (°C), Turb = Turbidity (NTU).

**Table 2.** The Principal Components Analysis (PCA) loadings for Cordillera Vilcanota and Cordillera Blanca for each environmental variable and the results of Pearson’s correlation tests between each PC axis and glacier cover (relationship with GCC%). Bold PCA loadings/correlation test results are those with moderate-strong (>0.3) correlations with PCA1 or PCA2 axes and bold p-values represent a significant relationship with GCC (p < 0.05) (Figure S1).

| Response variable | PCA1 | PCA2 |
| --- | --- | --- |
| Water temperature (°C) | **-0.315** | **-0.308** |
| Channel stability (1/Pfankuch) | -0.135 | -0.195 |
| Turbidity (NTU) | -0.090 | **0.630** |
| Conductivity ($\mu\text{S cm}^{-1}$) | **0.456** | -0.189 |
| pH | **0.338** | **-0.385** |
| Total nitrogen ($\text{mg L}^{-1}$) | **-0.367** | -0.280 |
| Total phosphorous ($\text{mg L}^{-1}$) | -0.240 | **0.304** |
| Dissolved organic carbon ($\text{mg L}^{-1}$) | **-0.312** | -0.294 |
| Dissolved inorganic carbon ($\text{mg L}^{-1}$) | **0.517** | -0.168 |
| Relationship with GCC% ($R^2$) | 0.058 | **0.292** |
| Relationship with GCC% (p-value) | 0.270 | **0.008** |

### Diatom biodiversity

#### α-diversity

Average taxonomic richness ranged from 2 taxa at 96% GCC in Vilcanota, to 31 taxa at 23% GCC in Blanca. Across both regions, taxonomic richness increased significantly as glacier cover decreased (Figure 3a & e; Table 3); however, the Cordillera Blanca region had a slight unimodal relationship, where richness was highest under low glacial influence. Whilst the unimodal relationship was not clear in the Vilcanota region, richness at sites with low glacial influence was also consistently high (3-18% GCC = 20-25 taxa) and similar to richness in Cordillera Blanca, showing rivers across regions with low levels of glacial influence can harbour similar or greater diatom diversity than rivers where no glacial influence is present. (Figure 3a & e).

**Figure 3.**
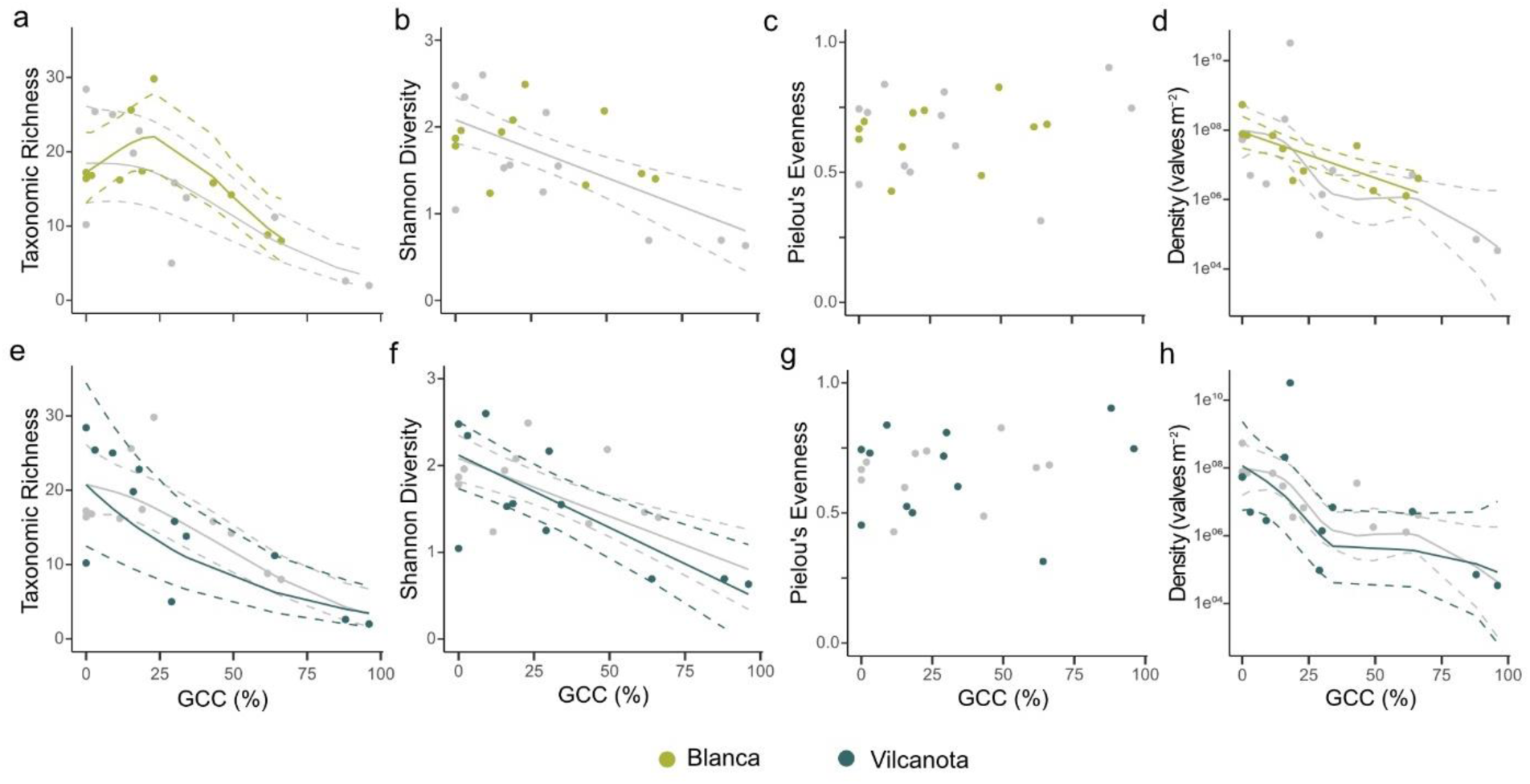
The relationships between glacier cover and α-diversity metrics of taxonomic richness (a, e), Shannon diversity index (b, f), Pielou’s evenness (c, g) and valve density (d, h). Green circles and lines on the upper row refer to sites and trends in Cordillera Blanca (a-d) and blue circles and lines on the lower row refer to Cordillera Vilcanota (e-h). Solid grey lines and grey circles represent the overall trends and data in both regions when combined and dashed lines are 95% confidence intervals. Trend lines are only included where there were significant relationships with GCC (p < 0.05) that were determined using GAMMs.

**Table 3.** Results of GAMMs and Mantel tests for relationships between α- and β-diversity metrics and glacier cover in the catchment (% GCC) in both regions (combined) and the Cordillera Vilcanota or the Cordillera Blanca regions. WS = within-site and BS = between-site. Bold and * p-values represent significant (p < 0.05) relationships.

| Response variable | Distribution | Combined |  | Vilcanota |  | Blanca |  |
| --- | --- | --- | --- | --- | --- | --- | --- |
|  |  | p-value | R<sup>2</sup> | p-value | R<sup>2</sup> | p-value | R<sup>2</sup> |
| Shannon diversity | Gaussian | **<0.001\*** | 36.1 | **0.005\*** | 55.5 | 0.339 | 0.8 |
| Taxonomic richness | Poisson | **<0.001\*** | 47.9 | **<0.001\*** | 53.6 | **0.027\*** | 61.6 |
| Pielou's evenness | Quasibinomial | 0.5 | 2.0 | 0.7 | 7.8 | 0.5 | 6.7 |
| Density (valves m<sup>-2</sup>) | Poisson | **<0.001\*** | 21.9 | **<0.001\*** | 39.7 | **0.003\*** | 14.2 |
| WS Total dissimilarity | Quasibinomial | **0.003\*** | 52.8 | **0.003\*** | 49.8 | 0.57 | 24.0 |
| WS Turnover | Quasibinomial | **<0.001\*** | 65.7 | **0.002\*** | 53.9 | 0.67 | 1.5 |
| WS Nestedness | Quasibinomial | 0.5 | 2.2 | 0.4 | 15.0 | 0.3 | 18.5 |
| BS Total dissimilarity |  |  |  | **0.04\*** | 14.1 | **0.05\*** | 28.0 |
| BS Turnover |  |  |  | 0.46 | <0.001 | 0.082 | 4.2 |
| BS Nestedness |  |  |  | **0.008\*** | 17.4 | 0.063 | 6.6 |

Shannon diversity increased significantly in the Vilcanota region with declining glacier cover but this trend was not apparent in the Blanca region. At comparable glacier cover levels between regions, Shannon diversity was at similar levels in both Blanca and Vilcanota; however, Blanca samples were not available from sites with very high GCC levels, where this trend with Shannon diversity became the most apparent in Vilcanota (Figure 3b & f; Table 3). No significant relationship between glacier cover and Pielou’s evenness was recorded in either region (Figure 3c & g; Table 3).

Average diatom valve density increased as glacier cover declined in both regions (Figure 3d & h; Table 3). Overall, density spanned seven orders of magnitude and ranged from 33,840 valves m^-2^ at 96% glacier cover to over 32 billion valves m^-2^ at 18% glacier cover. This maximum value was made up predominantly of *Encyonema* spp., and was more than 60 times higher than the next highest density, which was recorded at 0% glacier cover (540 million valves m^-2^).

#### β-diversity

Across both regions, within-site β-diversity was generally similar below 75% GCC but dissimilarity and turnover increased sharply above this level. Within regions, relationships between glacial influence and within-site β-diversity were only apparent in Cordillera Vilcanota, where total dissimilarity (Sørensen) and turnover (Simpson) declined with decreasing glacier cover. Turnover was an important influence on community structure within sites, most notably within high glacier cover sites. No significant relationship between within-site nestedness and glacier cover was observed in either region (Figure 4a-f; Table 3).

**Figure 4.**
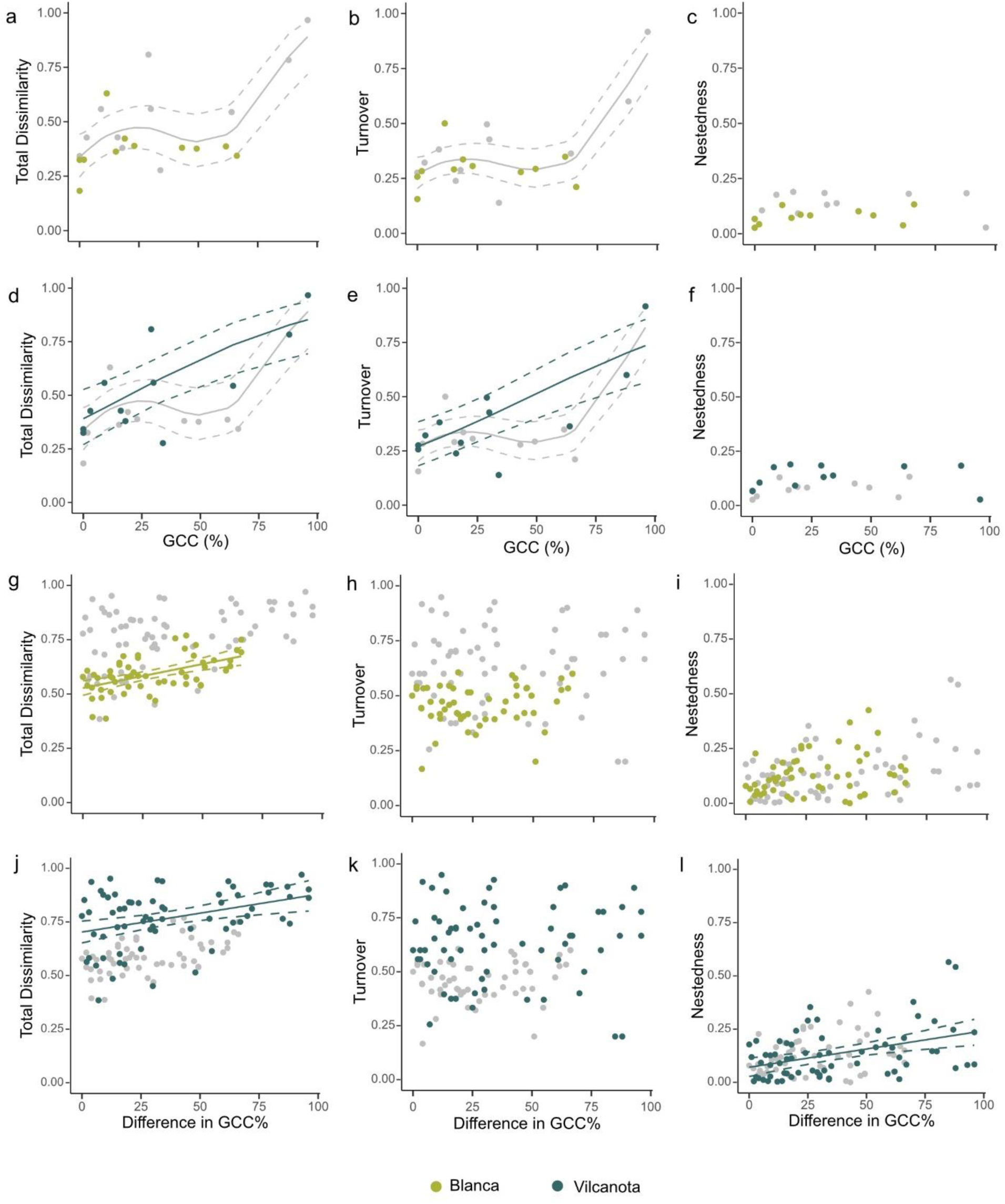
(a-f) β-diversity metrics for within-site total dissimilarity, turnover and nestedness in both Cordilleras Blanca (green; a-c) and Vilcanota (blue; d-f). (g-l) β-diversity metrics for between-site total dissimilarity, turnover and nestedness in Blanca (g-i) and in Vilcanota (j-l). Solid grey trend lines show significant relationships (p < 0.05) with glacier cover in the catchment (GCC) in both regions combined, green lines represent significant relationships in Blanca and blue lines in Vilcanota, determined using GAMMs (a-f) or Mantel tests (g-l). Dashed lines show 95% confidence intervals.

Between-site total dissimilarity was lowest in both regions when the difference in GCC between sites was also low, indicating sites with comparable glacial influence have similar species-abundance community composition. In Vilcanota, between-site nestedness also declined when glacial influence was more similar. Between-site species turnover and was not significantly related to glacier cover in either region (Figure 4g-l; Table 3).

### Community composition

In total, 130 diatom taxa from 32 genera were found in the Blanca region (0-66% GCC), compared to 176 taxa from 41 genera in Cordillera Vilcanota (0-96% GCC). Diatom assemblage composition varied regionally, with 119 species found exclusively in Vilcanota, 73 taxa identified only in Blanca and 57 taxa found in both regions. The most abundant diatom species in the Vilcanota region was *Encyonema minutum*, which made up 57% of the total valves and was most abundant at moderate levels of glacier cover. *Tabellaria flocculosa* was the most abundant species in the Blanca region, contributing to 33% of all identified valves and was found at all but one site. *E. minutum* was present at eight Blanca sites and *T. flocculosa* was found at just three Vilcanota sites and overall contributed to 1.25% and 0.002% of all valves, respectively.

In the Vilcanota region, *Reimeria sinuata* was recorded at the greatest number of sites (10 sites, representing 0-96% glacier cover) and in Cordillera Blanca, four species were recorded across all 11 sites (*Achnanthidium minutissimum*, *Brachysira arctoborealis*, *Frustulia* cf. *saxonica* and *Hannaea arcus*). Across both regions, 31 taxa were found exclusively at sites with 0% glacier cover, and 144 taxa identified only at sites with some glacier cover in the catchment (GCC > 0%). Ten taxa were found exclusively at sites with ≥ 62% cover, five in Cordillera Blanca and five in Vilcanota, with only one species found in both regions. *Gomphonema* sp. 1 was identified only at the site with 96% glacier cover (Table S2).

In the Blanca region, one species (*Encyonema* cf. *caespitosum*) was associated exclusively with high glacier cover sites in the indicator species analysis, but was also associated with low glacier cover sites in the Vilcanota region. In both regions, eight taxa were significantly related only to sites that had some glacial meltwater input, whilst only *Staurosira venter* was connected exclusively with no glacial influence. Many taxa were associated with several glacier cover levels; these were often common taxa found at multiple sites across both regions such as *Hannaea arcus* and *Encyonema silesiacum* (Table 4).

**Table 4.**
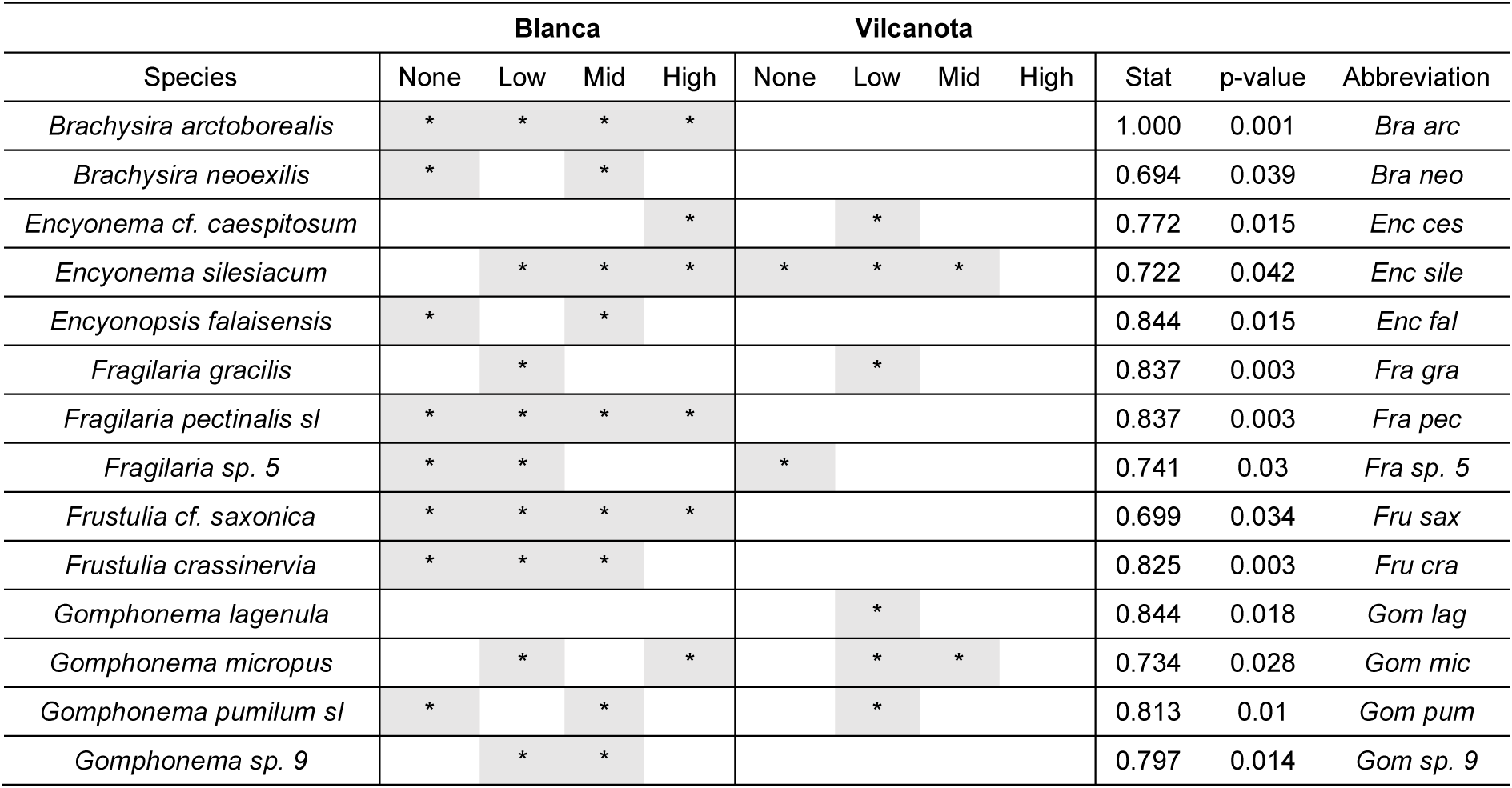
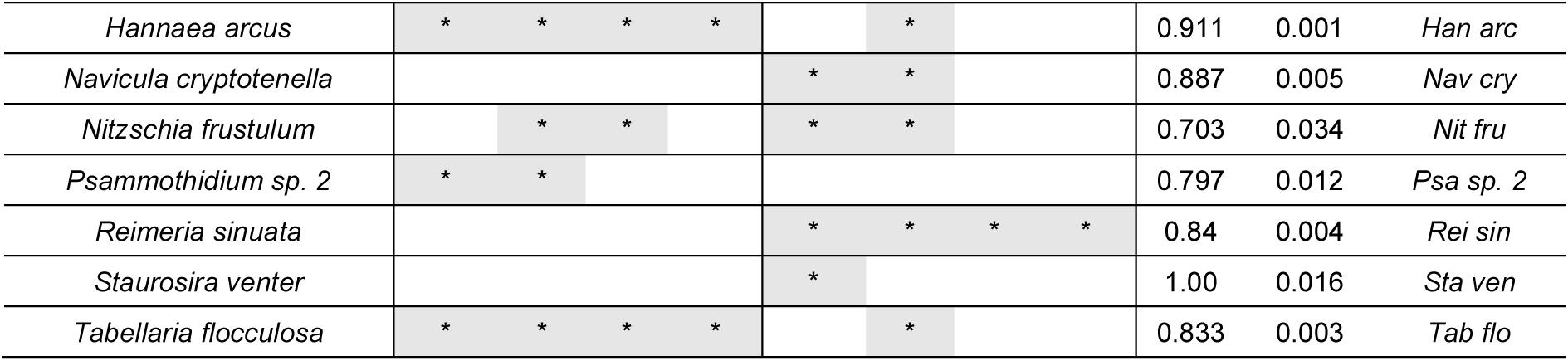
Diatom taxa that were associated significantly (p < 0.05; * and grey shading) with glacier cover in the catchment levels (None = 0%, Low = 1-25%, Mid = 26-50%, High = > 50%) in an indicator species analysis using presence-absence data. Abbreviations refer to taxa in Figure 5. “sl” = *sensu lato*.

The NMDS ordination showed the clustering of Blanca sites separately from most Vilcanota sites primarily along Axis 2. Axis 1 was associated with PC2 and GCC, representing physical river habitat characteristics, whilst axis 2 was correlated more closely with PC1, linked to water chemistry (Figure 5a; Table S3). Diatom taxa from the Vilcanota region were more widely spread along axis 1, and taxa found exclusively at high glacier cover sites, such as *Gomphonema* sp. 1, had high NMDS axis 1 scores (Table S4). Like the site clustering, taxa that were exclusive to the Blanca region had higher axis 2 scores, and species that were common across both regions tended to have NMDS scores closer to zero along both axes (Figure 5b). Taxa significantly associated with GCC levels in the indicator species analysis also tended to be taxa found at multiple sites, with mid-range NMDS scores (Figure 5b; Table 4).

**Figure 5.**
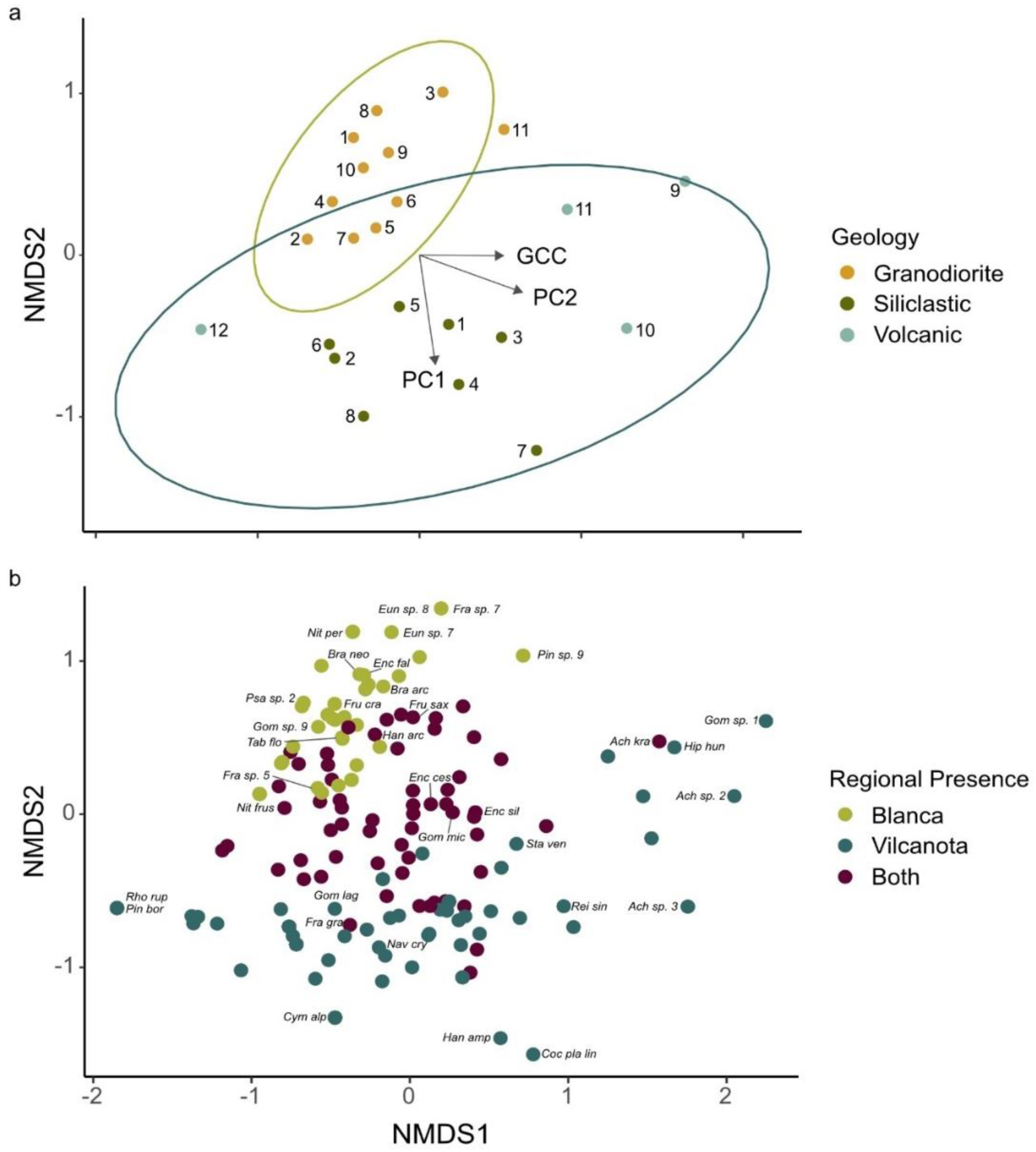
NMDS ordination plots of: (a) river sites coloured by underlying geology, ellipses which best enclose each region (green ellipse = Blanca, blue ellipse = Vilcanota), and significantly correlated PC axes and GCC. Numbers refer to sites within each region. (b) All species within the 2-dimensional ordination space, coloured based on presence in one or both regions. Abbreviated taxa names (full names in Tables 4 and S4) are included where they were either significantly associated with a glacier cover level in the indicator species analysis, or were at an extreme end of an NMDS axis (NMDS1 < - 1.5 or > 1.5; NMDS2 < -1.2 or >1.0). A Shepard Plot for the NMDS ordination (R^2^ = 0.962, stress = 0.195) indicated the accurate preservation of rank orders within the two-dimensional space, ensuring that the resulting NMDS configuration represented the original distribution of data within the dissimilarity matrix as closely as possible.

## Discussion

This inter-regional analysis is the first to show that declining glacier cover impacts assemblages of benthic diatoms in the Andes, and contributes new findings to an otherwise understudied group of organisms in glacier-fed rivers (Fell et al., 2017). Reductions in glacier cover affected diatom α-biodiversity consistently across both study regions, with similar increases in taxonomic richness and diatom abundance as glacier cover reduced, although low-mid levels of glacial influence often hosted elevated taxonomic richness. Whilst communities from Cordillera Blanca and Cordillera Vilcanota showed different responses in between- and within-site β-diversity components, across both regions most β-diversity metrics decreased as glacial influence declined, indicating that diatom assemblages as a whole can be expected to become more similar as glaciers retreat. Each region hosted distinct diatom assemblages, with just 23% of all 249 species shared among regions. Alarmingly, many of these regionally unique taxa were also found only at sites that had high glacial meltwater input, indicating that some diatom taxa will become threatened as glaciers continue to retreat, similar to findings from a number of European mountain rivers (Fell et al., 2018).

### α-diversity

Taxonomic richness and diatom valve density increased across both regions as glacier cover declined, supporting Hypothesis 1 and following similar trends seen at higher latitudes in Austria, Iceland, Italy and Canada (Hansen et al., 2006; Fell et al., 2018; Brahney et al., 2021; Bert et al., 2024). Slightly elevated α-diversity at low-mid levels of glacier cover compared to groundwater streams has also been recorded in diatom populations in Austrian rivers (Fell et al., 2018) and in macroinvertebrates in the same Cordillera Blanca rivers as this study (Palacios-Robles et al., 2024). In the southern hemisphere, increasing diatom abundance as glacier ice cover diminishes is supported by research from Ecuador that reported increased algal biomass (chlorophyll-*a*) in streams when glacier cover was low (Cauvy-Fraunié et al., 2016).

Increased diatom α-diversity with declining glacier cover, and higher diversity at low-mid levels of glacier influence, align with ecological stress theories such as the Intermittent Disturbance Hypothesis (Connell, 1978) and the harsh-benign concept (Menge and Sutherland, 1987), likely reflecting trade-offs in abiotic and biotic pressures structuring diatom assemblages. For example, turbidity, low water temperature and low channel stability were associated strongly with high glacier cover at Peruvian sites and these are key factors in shaping glacier-fed river biota (Milner et al., 2001; Fell et al., 2018; Bert et al., 2024). These conditions lead to frequent disturbance and restructuring of benthic communities (Brown and Milner, 2012) and supress biotic interactions such as competition and herbivory. As glacier cover decreases, changes in the timing and magnitude of flow events results in less stream bed disturbance and overall habitat harshness, leading to higher geomorphological stability (Cauvy-Fraunié et al., 2013; Khamis et al., 2016). However, lower abiotic stress in rivers with no meltwater contribution can induce biotic pressures through increased competition (Lange et al., 2011) or consumer pressure (Clitherow et al., 2013; Fell, 2019). Overall, α-diversity trends suggest that niche regulated assembly processes shape diatom assemblages across glacier river continuums in similar ways despite large differences in biogeography, but more research is needed to disentangle physical and chemical relative to biotic influences on diatom assemblages.

### β-diversity

Multiple β-diversity components decreased as glacier cover declined in both Cordilleras, indicating that diatom assemblages will become more similar both within and between streams as glaciers retreat, similar to findings from Europe (Fell et al., 2018). Within sites, divergent diatom assemblages at the patch (cobble) scale were strongest when glacier cover was > 66%. This variability in high glacial influence river sites may reflect time since the last disturbance of individual cobbles, due to low channel stability near glacier termini. Additionally, unmeasured flow hydrodynamics at the patch-scale are likely to be important for biofilm development (Krsmanovic et al., 2021) where flows are diurnally variable and often highly turbulent. These conditions are also important for invertebrate consumer presence at the cobble scale (Effenberger et al., 2006), synergistically influencing diatom colonisation and community development.

Between-site total dissimilarity decreased significantly in both regions as the difference in glacier cover amongst sites diminished, which is supported by studies of macroinvertebrate communities from the Blanca region (Palacios-Robles et al., 2024). As the range of glacially influenced rivers narrows in alpine systems, habitat heterogeneity between sites decreases (Milner et al., 2017), resulting in overall less niche space. As glaciers disappear completely, community composition may homogenise, threatening cold stenothermic or harsh-habitat specialists. This shift in diatom assemblage could also alter the grazing susceptibility of microalgae communities and have considerable implications for macroinvertebrates in these organic-matter limited ecosystems (Clitherow et al., 2013). In glacier fed rivers in Austria, between-site nestedness declined with reduced glacier cover differences (Fell et al., 2018), comparable to Cordillera Vilcanota, although turnover still contributed more highly to total dissimilarity. In support of Hypothesis 2, reductions in both within- and between-site β-diversity indicate that diatom assemblages in alpine regions may homogenise as glaciers retreat. However, the regional differences in β-diversity changes seen across multiple spatial scales emphasises the complexities in community responses to climate change and the need for further research from regions with diverse biogeographies.

### Diatom taxon and community responses

There was high regional diversity of diatom assemblages in the Peruvian Andes, similar to that found in other alpine systems (Cantonati, 2001; Falasco et al., 2012) with 119 species found exclusively in Vilcanota and 73 taxa identified only in the Blanca region. Whilst the overall lower diversity in Cordillera Blanca (130 in Blanca vs 176 in Vilcanota) may in part reflect the narrower range of glacier cover percentage sampled, NMDS axis 2 showed that water chemistry variables such as DIC and pH (associated with PC1), linked to different catchment geologies, were primary drivers of high regional diversity. Variation in catchment geologies in Vilcanota potentially resulted in a wider range of ecological niches for diatom species to occupy with factors such as ionic concentration and alkalinity likely to be key drivers in determining diatom species composition (Potapova and Charles, 2003). High conductivity was linked to elevated DIC, which is made up predominantly of carbonate (CO_3_ ^2-^) and bicarbonate (HCO ^-^) ions that diatoms can convert to CO for photosynthesis using external carbonic anhydrase enzymes (Knotts and Pinckney, 2019). CO_3_ ^2-^ and HCO ^-^ were key ions in shaping diatom assemblages in North America (Potapova and Charles, 2003). Taxa known to prefer high DIC (e.g. *Reimeria sinuata*) (Potapova and Charles, 2003), had low NMDS2 scores and were often found only in the Vilcanota region, characterised by sedimentary geologies more prone to weathering. Conversely, taxa such as *Brachysira arctoborealis*, known from soft-water bodies in North America (Wolfe and Kling, 2001) were characteristic of high NMDS2 scores and were only found in Cordillera Blanca. This shows that whilst meltwater input may influence overall diversity, individual species responses can also be dependent on individual catchment characteristics, highlighting a need for several catchments to be studied when trying to understand diatom responses to glacier retreat.

Diatom assemblages in the Peruvian Andes were dominated by a few generalist species, such as *Hannaea arcus*, *Encyonema minutum*, and *Tabellaria flocculosa.* Similar patterns of dominance were seen in the Canadian Rockies (Gesierich and Rott, 2012), Austrian and Italian Alps (Gesierich and Rott, 2004; Fell et al., 2018; Bert et al., 2024), Iceland (Hansen et al., 2006) and the Himalayas (Cantonati, 2001; Nautiyal et al., 2015). Many of these species were also significantly associated with several glacier cover groups, showing they can tolerate a wide range of glacial effects. Some of the same indicator species were identified in Italian alpine rivers (e.g. *H. arcus*), although the range in glacier cover scenarios was much lower (high glacial influence was considered as any sites > 11.5% GCC) (Bert et al., 2024). Low profile body forms, with strong attachment structures and low motility are characteristic of many dominant diatom taxa (e.g. *R. sinuata*) across alpine regions, including those identified in Peru. These traits help taxa withstand the high shear stress due to water flow and scour from suspended sediments in proglacial river environments (Biggs et al., 1998; Bert et al., 2024). Other common alpine taxa found in Peru, such as *T. flocculosa*, have larger cell sizes (Rimet and Bouchez, 2012), but their high reproductive rate may play a crucial role in persistence and recovery during frequent disturbances (Knudson, 1952; Biggs et al., 1998).

In support of Hypothesis 3, several taxa associated with no or low glacier cover (e.g. *Navicula cryptotenella* and *Gomphonema lagenula*) indicate that decreased GCC may allow a wider range of cell types to persist (Rimet and Bouchez, 2012; Zhang et al., 2018). These and many other taxa associated with low GCC have traits such as vertical attachment structures, large cell sizes (i.e. high profile taxa (Passy, 2007)) or moderate to high motility, potentially allowing better access to resources in established biofilms where competition may be more prevalent (Passy, 2007). Recent research in Italy found that prostrate ecological guilds (i.e. strongly attached to benthos) were dominant in streams with glacial meltwater input, and were replaced by motile and high profile taxa as glacier cover reduced (Bert et al., 2024). In Austria, a wider range of growth forms was also shown as glacier cover decreased, suggesting more mature biofilm establishment may be a consistent response of benthic microbiota across regions to glacier retreat (Fell et al., 2018). Diatom cell size and attachment structures can influence macroinvertebrate grazing ability (Feminella and Resh, 1991; Holomuzki and Biggs, 2006) so shifts in diatom functional diversity could impact higher trophic levels.

Across both regions, 38 taxa were found exclusively at sites with ≥ 25% glacier cover in the catchment, which is substantially higher than the six taxa identified at ≥ 28% GCC in Austria (Fell et al., 2018). This provides further support for Hypothesis 3, while also suggesting tropical diatom assemblages may be at greater risk of regional loss as glaciers retreat. Although only *Encyonema* cf. *caespitosum* had a significant association with high glacial input, the ten taxa found exclusively at ≥ 62% GCC are at risk of regional extinction as many glaciers in the Peruvian Andes are expected to decrease well below this high cover level (Rounce et al., 2023). Vulnerable taxa were mostly generalists known to span lowland and alpine environments such as *Gomphonema* spp. (Abarca et al., 2020; Kahlert, Maaria Karjalainen, et al., 2022) or those often found in cold-water environments such as *Eunotia* spp. (Castañeda Gómez et al., 2024). Whilst none of the taxa identified exclusively at sites with high glacier cover appear to be alpine or cold-water specialists (Round et al., 1990; Lange-Bertalot et al., 2017), the combination of regional endemicity, low motility of some taxa and habitat fragmentation caused by glacier ice loss may result in both geographic and genetic isolation of local populations, potentially leading to their decline as glaciers retreat. Additionally, restricted knowledge of diatom taxonomy in alpine regions of South America needs to be addressed to improve understanding of climate change effects on biodiversity, incorporating additional methods such as metabarcoding.

### Conclusions

This study illustrates clear changes in river diatom biodiversity with glacier retreat in South America. Diatom assemblages can be expected to become more similar both within and between river sites as glacier cover declines further into the future (Huss and Hock, 2018). Of particular importance is the potential threat to taxa that were found only within high glacier cover environments, of which few could be identified to known species. This suggests there may be heightened vulnerability to future deglaciation for some taxa but that further research is needed to fully quantify these threats.

As diatoms can form the primary food source for consumers in glacier fed rivers (Clitherow et al., 2013; Fell et al., 2017) and are highly important in supporting secondary productivity (Fellman et al., 2015), the altered biodiversity and abundance of diatom communities as glacier cover reduces may have cascading effects on higher trophic levels (Cauvy-Fraunié et al., 2016; Fell, 2019). The findings of this study correlate with diatom research from Austria (Fell et al., 2018), and other regions where glacier fed/ non-glacier fed systems have been studied (e.g. Canada (Brahney et al., 2021), Italy (Bert et al., 2024) and Iceland (Hansen et al., 2006)). Diatom assemblage responses to declining glacier cover may therefore be consistent across biogeographic regions. Global comparative studies using the same sampling and analysis methods would be useful to test this hypothesis directly. As human society in mountain regions relies heavily on glacial rivers for water supply (Mark et al., 2017), understanding the ecological consequences of glacier retreat on the primary producers of glacial river food webs is critical to predict the potential threat to water quality maintenance with future climate change.

## Acknowledgements

This research was financially supported by a Natural Environment Research Council (NERC) grant (NE/S013296/1) awarded to LEB and DQ, the Newton-Paulet Fund awarded to EL and KM, and the Leeds-York-Hull NERC Doctoral Training Partnership (DTP) Panorama studentship (NE/S007458/1), awarded to NRK. We thank Nilton Montoya and TodoVertical Cusco for assisting with fieldwork logistics and Leonore Debrov, Guadalupe Heredia, Hairo León, Edson Palacios-Robles and Sofia Rodriguez-Venturo for their assistance in sample collection. We also thank David Ashley for assisting in diatom sample preparation prior to identification.

## Author Contributions

NRK: Data curation; formal analysis; investigation; project administration; writing – original draft. DQ: funding acquisition; investigation; supervision; writing – review and editing. SCF: investigation; writing – review and editing. MK: investigation; writing – review and editing. KM: investigation; writing – review and editing; EL: funding acquisition; investigation; writing – review and editing. LEB: Conceptualisation; funding acquisition; investigation; supervision; writing – review and editing.

