## Supplementary Material for "Diatom biodiversity response to shrinking glaciers in the Peruvian Andes"

### Supplementary Figures and Tables

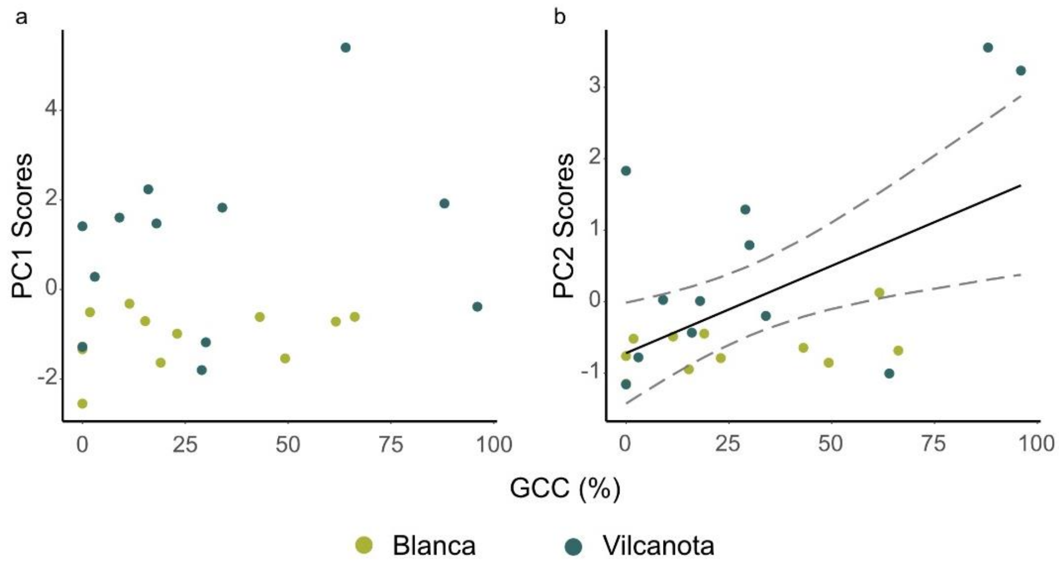


**Figure S1.** The relationship between glacier cover in the catchment (GCC %) and (a) PC1 and (b) PC2 scores in the Cordillera Blanca (green circles) and Cordillera Vilcanota regions (blue circles). Solid black lines are Pearson's correlation trend lines and dashed black lines are 95% confidence intervals.

**Table S1.** Physical and chemical river variables and the equipment used to measure each component at each site.

| Variable | Unit | Measurement tools |
|---|---|---|
| Glacier cover in the catchment (GCC) | % | Satellite data and QGIS tools |
| Electrical conductivity (EC) | μS $cm^{-1}$ | Hach HQ40D probe |
| pH | | Hach HQ40D probe |
| Water temperature | ºC | Hach HQ40D probe |
| Turbidity | NTU | Oakton T-100 turbidity probe |
| Channel stability | 1/Pfankuch | Pfankuch (1975) bottom component |
| Total nitrogen (EC) | mg $L^{-1}$ | Automatic wet chemistry analyser |
| Total phosphorus (TP) | mg $L^{-1}$ | Automatic wet chemistry analyser |
| Dissolved inorganic carbon (DIC) | mg $L^{-1}$ | Analytik Jena Multi NC2100 micro elemental analyser |
| Dissolved organic carbon (DOC) | mg $L^{-1}$ | Analytik Jena Multi NC2100 micro elemental analyser |

**Table S2.** Diatom taxa found exclusively at sites with mid-high glacier cover in the catchment (GCC %) in each region.

| GCC | Blanca | Vilcanota |
|---|---|---|
| High (> 50% +) | *Encyonema* cf. *cespitosum* | *Brachysira* sp. 3 |
| | *Eunotia* sp. 8 | *Frustulia vulgaris* |
| | *Fragilaria* sp. 7 | *Gomphonema* sp. 1 |
| | *Fragilaria* sp. 9 | *Gomphonema* sp. 6 |
| | *Pinnularia* sp. 9 | *Navicula gregaria* |
| Mid (26-50%) | *Encyonema* sp. 19 | *Achnanthes* sp. 1 |
| | *Eunotia silvahercynia* | *Achnanthes* sp. 2 |
| | *Eunotia* sp. 7 | *Achnanthes* sp. 3 |
| | *Fragilaria tenera* | *Aulacoseira* sp. 1 |
| | *Gomphonema* sp. 10 | *Brachysira* sp. 1 |
| | *Pinnularia* sp. 5 | *Caloneis* sp. 2 |
| | | *Caloneis* sp. 5 |
| | | *Encyonema* sp. 1 |
| | | *Eunotia* sp. 1 |
| | | *Hantzschia amphioxys* |
| | | *Hippodonta hungarica* |
| | | *Luticola* sp. 2 |
| | | *Odontidium hyemale* |
| | | *Pinnularia* cf. *borealis* |
| | | *Pinnularia* cf. *sinistra* |
| | | *Pinnularia* sp. 2 |
| | | *Pinnularia* sp. 3 |
| | | *Pinnularia* sp. 5 |
| | | *Planothidium* sp. 2 |
| | | *Psammothidium* cf. *helveticum* |
| | | *Pseudostaurosira* sp. 1 |
| | | *Rhopalodia* cf. *rupestris* |

**Table S3.** The correlations between PC axes and GCC with NMDS axes. Bold p-values represent significant correlations ($p < 0.05$).

| Variable | NMDS1 | NMDS2 | $R^2$ | p-value |
|---|---|---|---|---|
| PC1 | 0.145 | -0.990 | 0.47 | **0.003** |
| PC2 | 0.941 | -0.339 | 0.46 | **0.004** |
| GCC | 0.999 | -0.006 | 0.27 | **0.036** |

**Table S4.** Diatom taxa with NMDS scores that were either very high (NMDS1 > 1.5; NMDS2 > 1.1) or very low (NMDS1 < -1.5; NMDS2 < -1.1) on at least one axis. Higher NMDS1 scores were correlated to increased GCC and PC2 scores and higher NMDS2 scores were correlated with decreased PC1 scores (Table S3; Figure 5). Abbreviations refer to Figure 5.

| Taxa | NMDS1 | NMDS2 | Abbreviation |
|---|---|---|---|
| *Achnanthes sp. 2* | 2.05 | 0.12 | *Ach sp. 2* |
| *Achnanthes sp. 3* | 1.75 | -0.60 | *Ach sp. 3* |
| *Achnanthidium cf. kranzii* | 1.58 | 0.48 | *Ach kra* |
| *Cocconeis placentula var. lineata* | 0.78 | -1.57 | *Coc pla lin* |
| *Cymbella cf. alpestris* | -0.47 | -1.33 | *Cym alp* |
| *Eunotia sp. 7* | -0.12 | 1.19 | *Eun sp. 7* |
| *Eunotia sp. 8* | 0.20 | 1.34 | *Eun sp. 8* |
| *Fragilaria sp. 7* | 0.20 | 1.34 | *Fra sp. 7* |
| *Gomphonema sp. 1* | 2.25 | 0.61 | *Gom sp. 1* |
| *Hantzschia amphioxys* | 0.57 | -1.46 | *Han amp* |
| *Hippodonta hungarica* | 1.67 | 0.44 | *Hip hun* |
| *Nitzschia perminuta* | -0.36 | 1.19 | *Nit per* |
| *Pinnularia cf. borealis* | -1.85 | -0.61 | *Pin bor* |
| *Pinnularia sp. 9* | 0.72 | 1.04 | *Pin sp. 9* |
| *Rhopalodia cf. rupestris* | -1.85 | -0.61 | *Rho rup* |
